# Interaction of plant-derived metabolites and rhizobiome functions enhances drought stress tolerance

**DOI:** 10.1101/2024.12.06.627064

**Authors:** Anna Kazarina, Soumyadev Sarkar, Bryttan Adams, Leslie Rodela, Sophia Pogranichny, Eli Hartung, Loretta Johnson, Ari Jumpponen, Sonny T.M. Lee

**Affiliations:** Division of Biology, Kansas State University, Manhattan, Kansas, United States; Biodesign Center for Fundamental and Applied Microbiomics, Arizona State University, Tempe, Arizona, United States; Department of Biology, Texas State University, San Marcos, United States of America

**Keywords:** microbiome, plant-host microbe interactions, shotgun metagenomics, metabolomics

## Abstract

**Background:** Plants evolved alongside microbes, enabling plants to better cope with abiotic and biotic stresses. The interactions between plant roots and local soil microbes are critical for environmental adaptation and plant health. Plants actively regulate the microbial community composition in their rhizospheres to recruit specific microorganisms that enhance their fitness in the ecosystem they inhabit. This study builds on prior research suggesting that plants have a “home field advantage” in recruiting microbes unique in their home environment, reflecting mutual recognition and the targeted recruitment of microbes.

**Results:** Using gene- and genome-centric approaches, we assessed the functional potential of root-associated microbes and profiled the host metabolites to uncover the metabolic outputs potentially regulating host-microbe interactions. Our results showed that plants adapted to drier environments experience less stress, producing fewer stress-related metabolites and impacting the recruitment of microbes with genes linked to stress relief pathways. In particular, plant-derived trimethyllysine was highly associated with microbial populations capable of improving nutrient uptake, producing plant growth-promoting compounds, and modulating stress responses.

**Conclusion:** This study highlights the critical interplay between host exudates and microbial substrate uptake as the primary mechanism of rhizosphere assembly. We demonstrate that plants actively produce metabolites to recruit microbial populations with the functional potential to enhance hosts’ ability to thrive in a stressful environment. This research provides insights into the mechanisms of plant-microbe communication, rhizosphere recruitment, and the complex interplay of plant-microbe interactions. Furthermore, it highlights promising avenues for manipulating rhizosphere microbiomes to support conservation agriculture in the face of climate change.

## Background

The “home-field advantage” hypothesis in plant-soil microbe interactions posits that plants thrive better when growing in an environment containing microbial communities that have co-evolved with or co-adapted to over time [1–5]. Our previous work has shown that plant hosts perform better at recruiting microbes unique to their “home” environments, suggesting a mutual recognition between the plant host and local soil microbes [6]. These results also suggest that plants originating from more stressful conditions, such as saline or drought-prone environments, selectively recruit specialized beneficial microbes for their functional potential to reduce the host’s stress [7–10]. Understanding how local soil microbes confer functional advantages to plant hosts is essential for fully grasping the dynamics of interactions between plant hosts and microbes as well as their implications for plant health and resilience.

Numerous studies have demonstrated the effects of specific bacterial taxa in influencing plant health and growth [11–15]. For example, the bacterial genera *Pseudomonas* and *Rhizobium* are known for their nitrogen-fixing capabilities, enhancing the host’s nutrient availability [12, 16]. Additionally, *Bacillus* and *Burkholderia* can contribute to phosphorus solubilization, further supporting plant growth and resilience [17–19]. While studies on the specific taxa reveal valuable insights into the plant-microbiome interactions, we suggest that the rhizosphere recruitment is not based on the attraction of specific taxa, but rather on the functional potential of microbes [20]. Additionally, while studies exploring system dynamics rely on the taxonomic assignment of the sequencing data to evaluate this interaction, we know little about how rare, low-detection, or uncultured microbial taxa can impact the plant host or other microbes.

Plant-soil microbe recognition and communication occur through complex nutrient and metabolite exchanges, where plant-released exudates can be utilized by rhizosphere microbial communities [21, 22]. The rhizosphere is surrounded by a soil matrix where the plant roots continuously produce and secrete a diverse array of organic metabolites, which vary in chemical compositions and are mostly controlled by the host genetics [23], host developmental stage [24], and many abiotic and biotic factors [20, 22, 25–28]. Plant metabolites also serve as a nutrient source and signaling molecules to facilitate the repelling of pathogens and the recruitment of beneficial soil microbes. Thus, host nutrient acquisition, pathogen resistance, and overall are enhanced [22, 29]. The rhizosphere microbiome (rhizobiome) consistently differs from the soil microbiome, primarily because of the selective influence of plant root exudates [20, 30]. Some metabolites selectively the growth of specific microbes and differentiate between beneficial and pathogenic microorganisms [31, 32]. For example, plants can exudate compounds for various purposes: aromatic organic acids (nicotinic, shikimic, salicylic, cinnamic, and indole-3-acetic) as energy sources for specific microbial populations; signaling molecules (phytohormones, specialized metabolites, volatile organic compounds, and peptides) or “cry for help” compounds like terpenes in response to biotic stress, attracting beneficial rhizosphere microbes [22, 31, 33–35]. This complex plant host-microbe interaction can result in variations of microbial composition even among individuals of the same plant species. While many studies have used the local adaptation of plant ecotypes to further our understanding of the plant-rhizobiome interactions [27, 36–38], little information exists on the impact of ecotypic differences in plant-produced metabolic profiles on the rhizosphere microbial community and functions.

Abiotic factors can drive shifts in the plant-associated microbial communities and functions. For example, sulfate-reducing bacteria (SRB) are abundant in waterlogged soils because of their anaerobic respiration abilities [39, 40], whereas others like Actinobacteria dominate drought-tolerant communities [41–43]. The plant-host adaptation to local environments likely also resulted in co-adapted microorganisms which also undergo reciprocal evolutionary changes that select for specific associations [44–46]. Like their plant host, microbes face environmental stresses, resulting in microbial populations with the specific genetic or functional potential to thrive under unfavorable conditions [47–50]. This may provide plant hosts with complementary benefits and the mutual adaptation can enhance the resilience and productivity of both plants and their associated microbial communities [51, 52]. This dynamic process influences the genomes, behaviors, and ecological roles of the hosts and microbes, fostering interdependent and often specialized relationships that account for plant-microbiota interactions’ diversity, specificity, and stability [53, 54]. Although this is a crucial aspect of the plant-host-microbe interaction, information on the chemical exchange between the plant host and its associated microbes remains limited.

This study focuses on the interaction between the plant host exudates and associated rhizobiome functional capabilities, within the framework of the plant as a holobiont—a complex, integrated system composed of the plant host and its microbial communities. We aim to deepen the understanding of the mechanisms underlying plant-microbe and microbe-microbe recognition, communication, and recruitment. Building on our previous work, we (1) investigated the functional potential of the recruited rhizosphere microbes through comprehensive gene-centric and genome-centric analyses of rhizosphere communities; and (2) adopted a holistic approach in studying plant holobiont systems by integrating microbial data with plant-host metabolic profiles, attempting to provide providing a comprehensive view of the complex interplay between local microbial communities and the plant host. We hypothesized that plants adapted to drier environments would experience less stress, resulting in fewer stress-related metabolites, which in turn, will impact the recruitment of the microbes with the functional potential of providing host stress relief. Our work improves the understanding of factors that influence the plant host’s rhizobiome recruitment and provides insight into host ecotype responses in the face of a changing climate.

## Results and Discussion

In this study, we used both community gene-centric and genome-resolved metagenomics, paired with host metabolomics, to elucidate how plant ecotypic characteristics and differences in mean annual precipitation (MAP) influence plant rhizobiome interactions in the dominant tallgrass prairie grass *Andropogon gerardii* (Figure 1).

**Figure 1.**
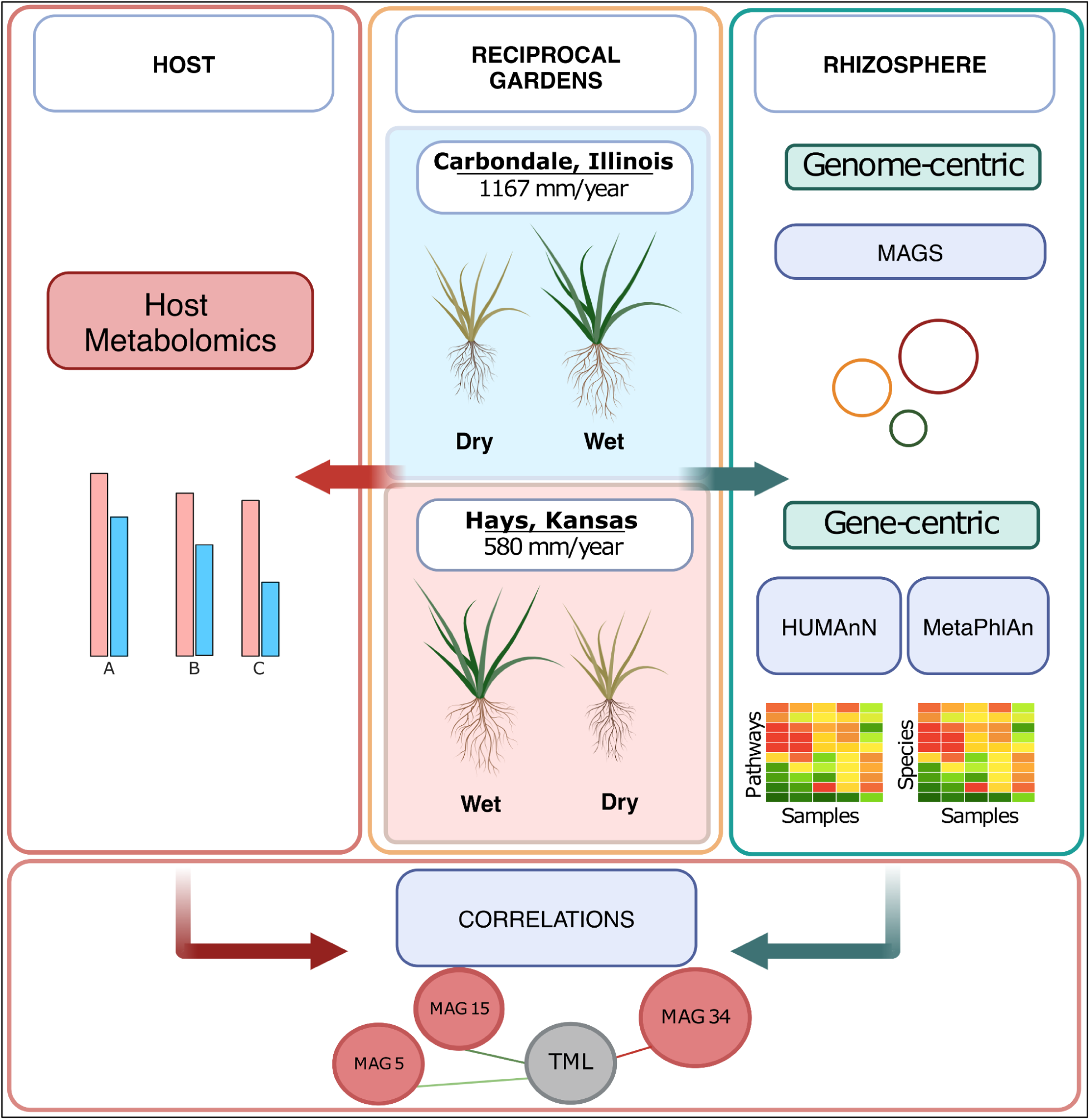
The schematics of the experimental design highlighting the flow of the project. Briefly, we collected rhizosphere samples from reciprocal gardens in Carbondale, Illinois, and Hays, Kansas, growing Dry and Wet regional *A. gerardii* ecotypes. We conducted shotgun metagenome sequencing to evaluate the microbial functional potential of these rhizosphere samples using gene- and genome-centric approaches. First, we performed gene-centric taxonomic community profiling (MetaPhlAn) and metabolic pathways profiling (HUMAnN). Next, we assembled 34 non-redundant Metagenome-Assembled Genomes (MAGs) and performed genome-centric profiling of the MAGs. To assess the host metabolic output, we performed plant-host metabolites profiling. Finally, we conducted a correlation analysis to identify key omics variables across the datasets.

### Distinct metabolic profiles between the Dry and Wet ecotypes reflect the Dry ecotype’s adaptation to the “home” environment

We detected 1,191 plant metabolites across all 24 rhizosphere samples from Hays and Carbondale (Figure 2, Table S2). Overall, metabolites were classified as amino acids (17.72%), others (13.85%), phenolic acid (12.68%), lipids (12.68%), alkaloids (9.49%), organic acids (8.73%), flavonoids (6.88%), terpenoids (5.46%) nucleotide and derivatives (5.04%), lignans and coumanins (4.70%), quinones (0.92%) and tannins (<0.01%) (Figure 2A, Table S2).

**Figure 2.**
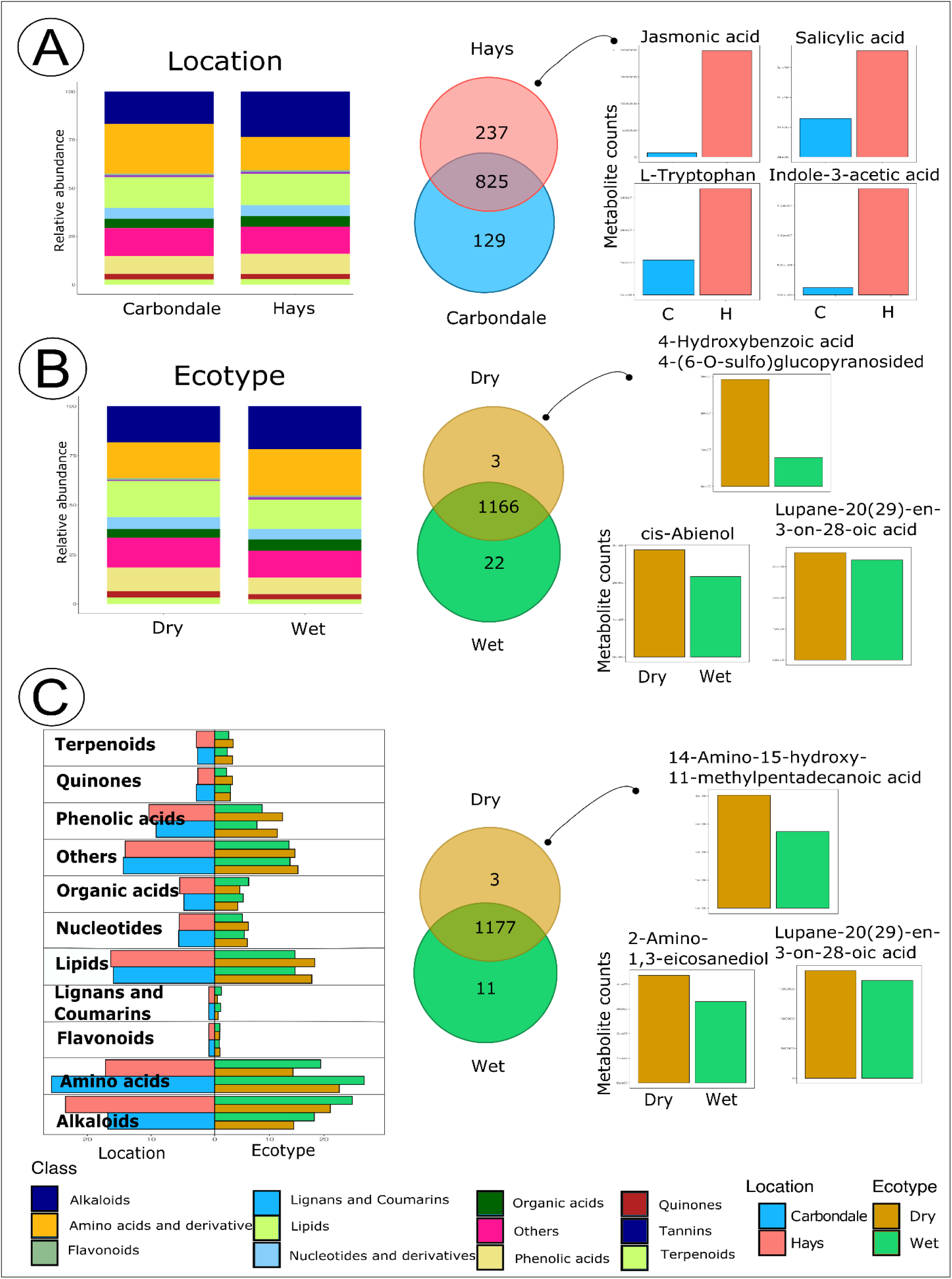
Plant hosts displayed different metabolic profiles across ecotypes and reciprocal garden locations. (A) The relative abundance of the plant host metabolites within Hays and Carbondale. The Venn diagram with the number of metabolites significantly different (not overlapped; p<0.05) across two locations and bar plots with four metabolites (shortlisted metabolites) significantly higher in Hays (p<0.05). (B) The relative abundance of the plant host metabolites within Dry and Wet ecotypes. The Venn diagram with the number of metabolites significantly different (not overlapped; p<0.05) across two ecotypes and three metabolites that were significantly higher in the Dry ecotype. (C) The two-sided bar plot represents the split of the metabolic classes across two ecotypes (Dry, Wet) between the two locations (Hays, Carbondale). The Venn diagram with the number of metabolites significantly different (not overlapped; p<0.05) between the two ecotypes in Hays and the three metabolites that were significantly higher in the Dry ecotype in Hays.

We observed subtle metabolome differences between the Dry and Wet *A. gerardii* ecotypes (Figure 2B, Table S2). In the Dry ecotype, the predominant metabolite classes were alkaloids (18.18%), amino acids (18.19%), and lipids (18.11%). The Wet ecotype had a higher proportion of alkaloids (21.73%), amino acids (23.43%), and a lower proportion of lipids (14.73%) than the Dry ecotype (Figure 2B, Table S2). The metabolites associated with the Dry ecotype included Lupane-20(29)-en-3-on-28-oic and 4-Hydroxybenzoic acid 4-(6-O-sulfo)glucopyranoside compounds with potential antimicrobial antioxidant properties [55, 56], and cis-Abienol - antimicrobial metabolite [57] that could enhance plant stress tolerance and defense mechanisms.

The observed 366 metabolites that varied between locations (ANOVA, Hays: n=237; Carbondale: n=129) (Figure 2A, Table S2) provided further insights into the ecotypic host-rhizobiome interactions that may enhance plant resilience in drier regions. We identified a diverse array of metabolites that were differentially more abundant in the drier Hays site than in the wetter Carbondale site. These metabolites were associated with plant-microbe communication (e.g., indole and indole-3-acetic acid (IAA)), as well as compounds involved in plant immune responses, such as jasmonic acid (JA) and salicylic acid (SA), L-Tryptophan. Only 25 metabolites differed between the *A. gerardii* ecotypes (Dry: n=3; Wet: n=22) (Figure 2C, Table S2). In pairwise comparisons between the Dry and Wet ecotypes in Hays, we identified 14 metabolites that differed between them (Figure 2C). Only three of these (14-Amino-15-hydroxy-11-methylpentadecanoic acid, 2-Amino-1,3-eicosanediol, and Lupane-20(29)-en-3-on-28-oic acid) were elevated in the Dry ecotype (Table S2). Notably, Lupane-20(29)-en-3-on-28-oic acid and 14-Amino-15-hydroxy-11-methylpentadecanoic acid can exhibit antimicrobial properties [58], a finding consistent with the observed antimicrobial metabolites in our overall ecotype analysis across the two sites. 2-Amino-1,3-eicosanediol is involved in the synthesis of Eicosanedioic acid (EDA), which can function as a building block for the construction of complex lipids essential for resilience in drier environments [59].

Our metabolic profiling indicated that ecotypes adapted to their environment in their responses to the drier conditions. We further demonstrated that Dry and Wet ecotypes may respond differently in the context of plant-microbe interactions, even within the same location. This crucial interaction between the ecotypes and their associated rhizobiome communities may be essential to understanding plant host resilience to drought. As a result, we asked-*Which microbial pathways could be crucial in the Dry ecotype-microbe interaction and could enhance the plant host resilience in a drier environment?*

### Microbial communities differed in functional potential and their impact on the plant host

After obtaining insights from the plant metabolic profiles, we focused on profiling the rhizosphere microbial diversity and pathways within the entire community to answer whether any specific taxa (Table S3) or gene functions (Table S4) were associated with the ecotypes or locations. Shotgun sequencing of the samples (n=24) resulted in approximately 1.8 x 10^9 sequences with an average of ∼ 77 x 10^6 ± 12 x 10^6 reads per metagenome (Table S5). We used MetaPhlAn and showed that of 2,538 bacterial taxa (Table S3), only 1,557 were classified to the genus level. We further showed that the rhizosphere microbial communities differed between the locations and ecotypes (Figure 3A). With the insights from plant metabolomes and rhizobiome composition differences [6], we focused on elucidating ecotypic rhizobiome differences in Hays, with a specific question: what microbes and pathways might contribute to the home-field advantage hypothesis and enhance Dry ecotype resilience in the drier environment in Hays [1, 2, 5]. In Hays, only four microbes differed between the Dry and Wet ecotypes (Figure 3B, ANOVA: p< 0.001; Hays: Wet vs Dry: p<0.025). Notably, we observed a higher abundance of *Kribbella speibonae* in the Dry ecotype in this drier site, suggesting its importance in enhancing host growth in the drier environment. *Kribbella speibonae* can catalyze 3-Phosphonatooxypyruvate to phosphoserine, which is crucial in the amino acid biosynthesis, including serine, cysteine, and glycine, all vital for plant growth and development [60–62]. In contrast, the Wet ecotype had a higher abundance of Kribbella sp. VKM Ac 2572 (n=2) (Figure 3B). The observed differences in the abundance of these bacteria, even at the genus level, likely indicate potential variations in their functional and ecological roles. This aligns with our observation that rhizosphere recruitment is driven not by the attraction of specific taxa but by the functional potential of the recruited microbes [20, 63, 64]

**Figure 3.**
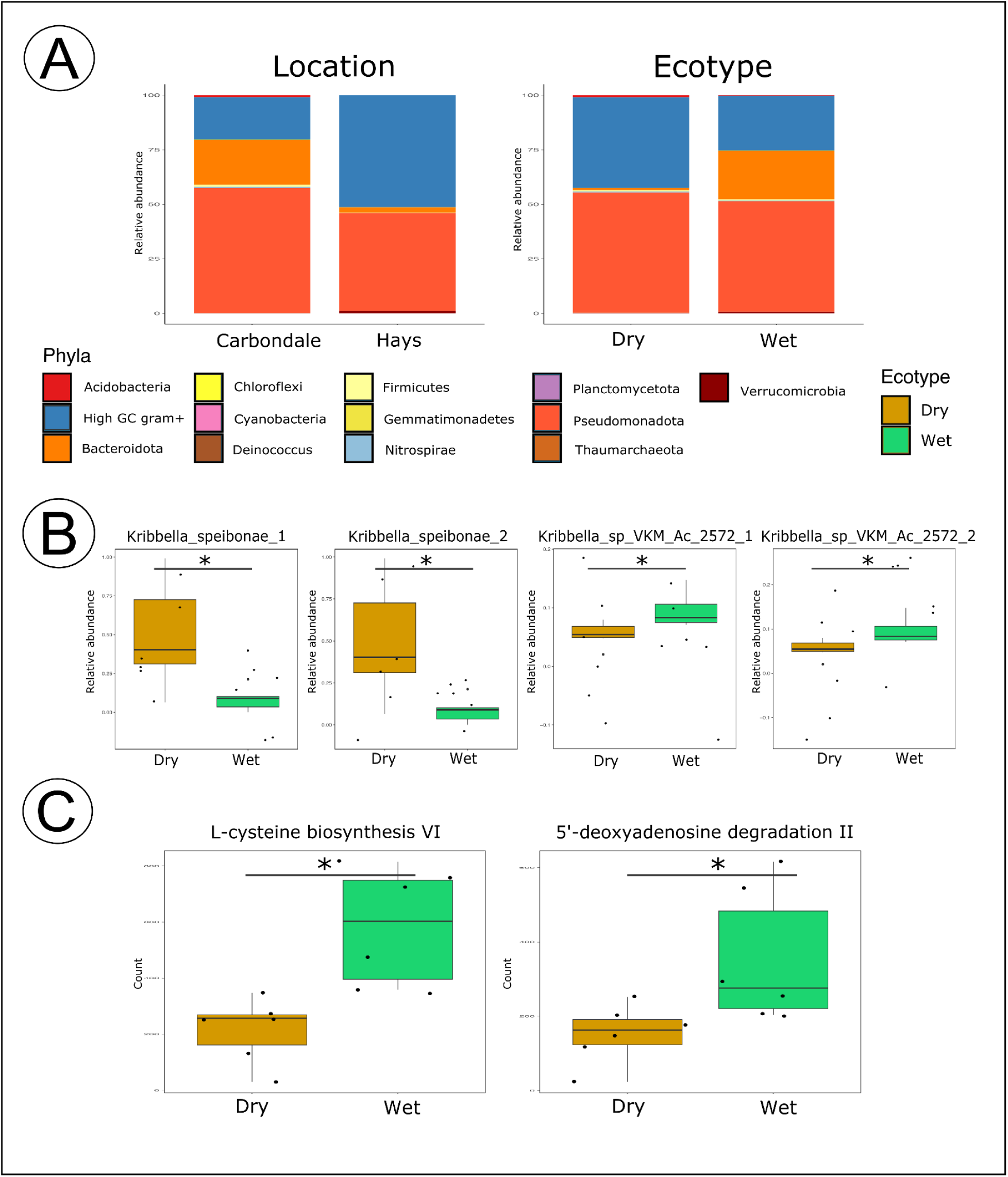
The gene-centric profiling of potential bacteria taxa (MetaPhlAn) and metabolic pathways (HUMAnN) corroborated the home-field advantage hypothesis, showing the Dry ecotype displaying resilience in the drier environment of Hays. (A) Stacked bars with the relative abundance of the taxonomic phyla within two locations and two ecotypes. (B) The abundance of four *Kribella* species that significantly differed between the Dry and Wet ecotypes in Hays. (C) Two metabolic pathways thant significantly differed between the Dry and Wet ecotypes in Hays, with the higher count associated with the Wet ecotype. Significant p-values marked by asterisks * represent p < 0.05.

We next performed a functional analysis (HUMAnN) of metabolic characteristics potentially contributing to the overall functioning of the rhizobiome across locations and ecotypes (Table S4). Although we identified a total of 548 pathways across the metagenome samples (Figure 3C), only two - 5’-deoxyadenosine degradation II and L-cysteine biosynthesis VI (from L-methionine) - differed significantly (ANOVA: p< 0.013; Hays: Wet vs Dry: p<0.026) between the Dry and Wet ecotypes and both were highly associated with Wet ecotype (Figure 3C). The predominance of L-cysteine biosynthesis in the Wet ecotype-associated microbes corroborated our home-field hypothesis that the Wet ecotype would not be as efficient in recruiting organisms that aid in environmental resilience as its Dry ecotypic counterparts. Microbe-associated L-cysteine biosynthesis facilitates the L-cysteine synthesis from L-methionine in a four-step process, in which the first two steps occur in the methionine cycle. At the same time, the latter two are part of the transsulfuration pathway. The L-cysteine synthesis by Wet ecotype associated microbes could act as an antioxidant and be crucial in the production of glutathione, an essential molecule for mitigating oxidative stress for its associated plant host [65–67]. More interestingly, we also noted that the first step of L-cysteine biosynthesis pathways produces S-adenosyl-L-methionine (SAM), a critical methyl donor in many biochemical reactions. Radical SAM enzymes convert SAM into a toxic by-product - 5’-deoxyadenosine. The differentially higher abundance of the 5’-deoxyadenosine degradation pathway in the Wet ecotype suggested an upregulation of various stress-related biochemical pathways [68]. These pathways might include the synthesis of “stress ethylene,” potentially triggered by environmental challenges such as drought and/or salinity stress in Hays. Our findings on microbial SAM-stress-associated pathway interactions suggest the potential exchange between the Wet ecotypic plant host and its rhizobiome under a drier, non-native environment (Hays). This resulted in increased plant host stress signals and the corresponding byproducts. The production of plant-host stress-associated products might have attracted or activated pathways in the rhizosphere microorganisms [69–72], enhancing their potential to alleviate stress.

### Distinguishable microbial populations and pathways were associated with ecotype and MAP

While community-level metagenome analyses provided insights into which microbial pathways might influence the plant-rhizobiome interaction, we also aimed to better understand which microbial populations and functions could impact the plant host resilience. To this end, we resolved a total of 34 non-redundant MAGs in our genome-centric metagenomic analysis (Figure 4, Table S6). These MAGs were assigned to the following phyla: Patescibacteria (6), Pseudomonadota (5), Desulfobacterota (2), Nitrospirota (2), Verrucomicrobiota (2), Bacteroidota (2), High GC gram+ (2), Acidobacteriota (1), Planctomycetota (1) all archaea were assigned to Thermoproteota (7), and four MAGs remained unassigned (Figure 4A). We observed that MAP was the main driver of the differences in rhizobiome microbial composition [73–75] (Figure 4B). Among the 34 resolved MAGs, we observed a higher detection of the bacterial phyla Actinobacteria (2), Acidobacteria (1), and Desulfobacterota (2) in the drier Hays location. While we anticipated that the drier conditions in Hays would favor taxa such as Actinobacteria and Acidobacteria, we were surprised to detect Desulfobacterota in this drier site, as Desulfobacterota typically thrive in water-saturated environments [76, 77]. This finding suggests that the potential plant host selection of rhizobiome microbes may be more driven by the microbial functional potential than their simple taxon affinities. In this vein, the determining factors could include microbial functions that sustain plant hosts’ activity and/or mechanisms that permit the host to thrive in unfavorable environments. In wetter Carbondale location, we observed high detection of MAGs annotated to bacterial phyla Pseudomonadota (4), Patescibacteria (4), Bacteroidota (2), and archaeal phylum Nitrospirota. Similar to Hays, the presence of these microbial groups aligned with their ecological roles in nitrification and organic matter decomposition in wetter soil environments.

**Figure 4.**
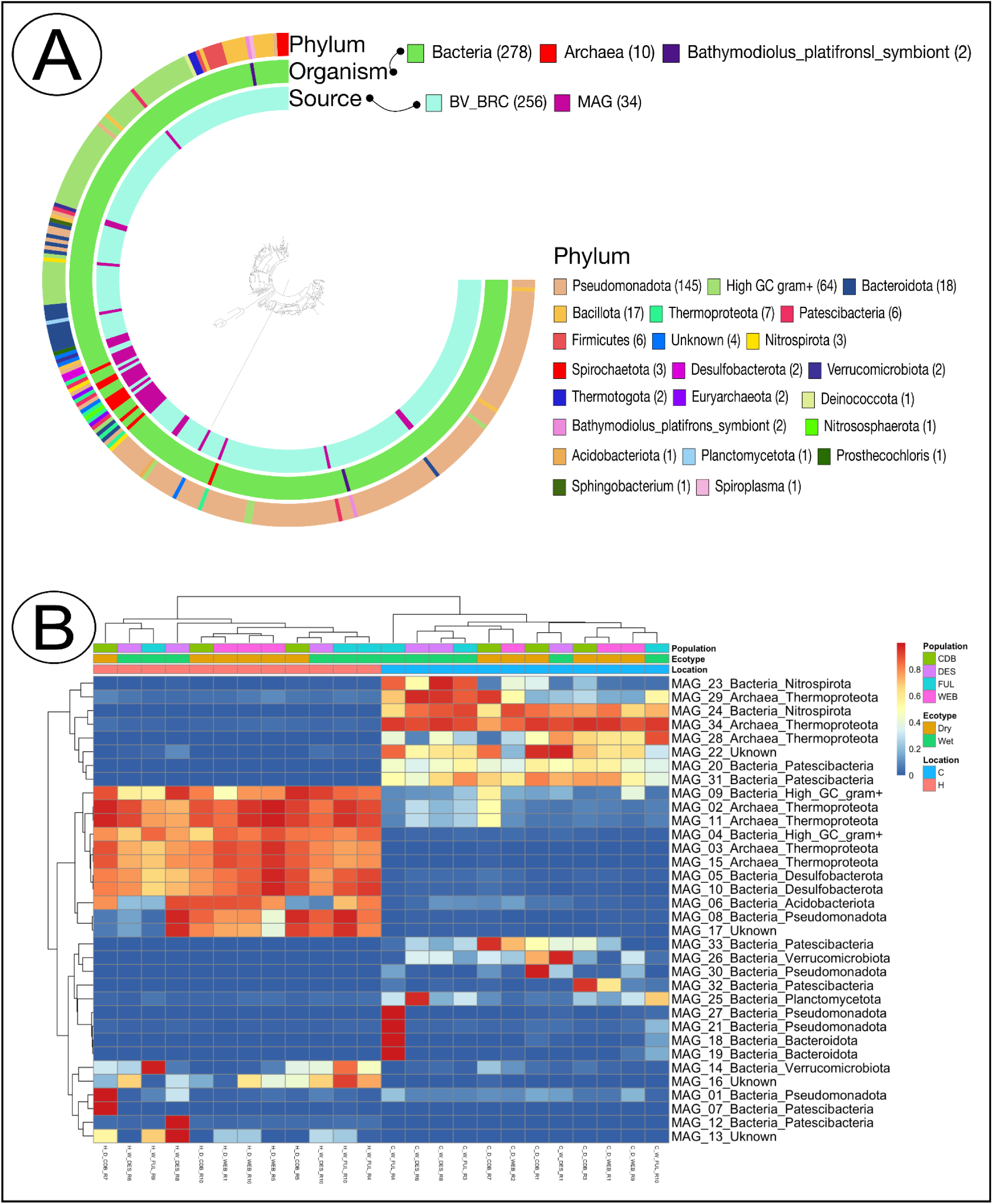
The phylogenomic placement of the MAGs and the detection of MAGS across collected rhizosphere samples. (A) Phylogenomic tree based on alignment of six bacterial ribosomal genes: Ribosomal_L1, Ribosomal_L2, Ribosomal_L3, Ribosomal_L4, Ribosomal_L5, Ribosomal_L6. The tree included 34 MAGs and 11 unique closest strains (a total of 324 bacterial strains) to identify relationships and the confirmation of the identity of MAGs. (B) Clustered heatmap with rows corresponding to 34 MAGs and columns corresponding to rhizosphere samples. The heatmap was clustered by rows and columns to observe the highest detection similarity within samples. The precipitation (location) was the main driver of changes in the detection of the MAGs.

We next investigated the functional potential of these 34 MAGs to understand 1) how the MAGs that were more commonly detected in Hays may have adapted to the drier conditions and 2) the implicated microbe-microbe and host-microbe communication functions in these microbial populations.

### Microbial survival in the environment

Our previous work hypothesized that specific microbial populations could provide beneficial functions to the plant host under stressful environments [6]. This dependency on the microbial function arises from the unfavorable environmental conditions that the plant host might experience [23, 50, 78–80]. As such, the plant-host’s stress signaling molecules and microbial functional requirements in the dry Hays environment might differ from those in wetter Carbondale. Therefore, the specialization of the microbial functional potential and their ability to survive in stressful environments go hand in hand [41–43, 64, 81], with bacterial survival relying heavily on their ability to sense and respond to environmental changes [82]. Here, we observed several mechanisms that microbial populations highly detected in Hays might utilize to survive in the drier environment (Figure 5, Table S7). For example, MAGs 1, 27, and 30 (all assigned to phylum Pseudomonadota) had all necessary EnvZ and OmpR regulon genes used for regulating the expression of outer membrane porins in response to osmotic stress [83, 84]. In addition, these MAGs also had an active BarA-UvrY(SirA) two-component regulatory system involved in bacterial sensing and responding to changes in their environment. Other highly detected microbial populations in this study, including MAGs 8 (Pseudomonadota), 15 (Thermoproteota), 17 (unassigned), 27 (Pseudomonadota), and 30 (Pseudomonadota) had the genomic potential to produce both (GlrK and GlrR) proteins of the GlrKR two-component system responsible for regulating gene expression in response to environmental stimuli [85, 86]. The survival capacity of these microbial populations is the first of many requirements for enhanced plant host resilience to abiotic stress due to plant-microbe interactions. With the identification of numerous microbial survival mechanisms in our resolved MAGs, we next investigated the functional potential of host-microbe and microbe-microbe communication capabilities.

**Figure 5.**
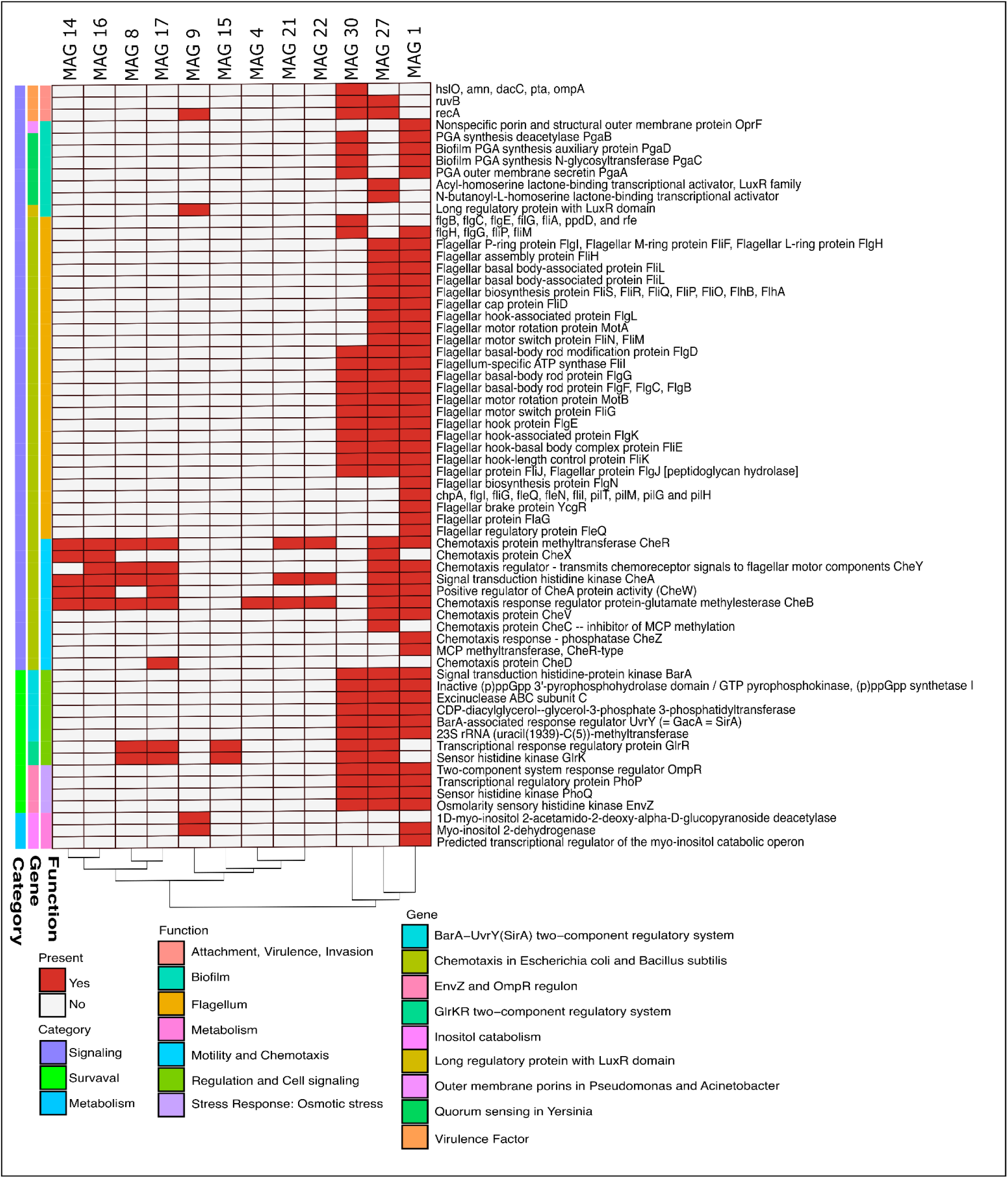
Genes/subsystems (subset) presence (red) and absence (grey) in selected MAGs. Rows correspond to the subset of genes/subsystems related to microbial mechanisms highly detected in Hays potentially unitized to survive in the drier environment, and columns correspond to the MAGs these genes/subsystems were observed in.

### Host-microbe and microbe-microbe signaling

To provide a direct beneficial function to the plant host, soil bacteria must become a member of the plant-host rhizosphere. We considered multiple levels of interactions when describing the possibility of plant-microbe communication. We explored the genomic potentials of the MAGs in their ability to perceive and interpret plant-host signals, their physical capacity to navigate toward these signals, rhizoplane biofilm formation, and their ability to bypass the plant immune responses to integrate successfully into the rhizosphere or endophyte community. *First*, we investigated the chemotaxis signaling, which is critical in host signal recognition (Figure 5, Table S7). This process begins with detecting the signals by transmembrane chemotaxis receptors, specifically methyl-accepting chemotaxis proteins (MCPs) [87, 88]. We observed that MAGs 1 (Pseudomonadota), 14 (Verrucomicrobiota), 16 (unassigned), 17 (unassigned), and 27 (Pseudomonadota) possessed all the necessary genes for CheA histidine kinase and the coupling protein CheW, which are required to form a ternary complex with MCPs for effective signal transduction [89, 90]. This interaction forms a two-component system with CheY, where phosphorylated CheY (observed in MAGs 1 (Pseudomonadota), 2 (Thermoproteota), 8 (Pseudomonadota), 16 (unassigned), and 17 (unassigned)) activates motor proteins responsible for bacterial motility [91, 92]. Additionally, we observed that MAGs 1 (Pseudomonadota) and 9 (High GC gram+) contained genes necessary for catabolizing the plant-exuded myo-inositol (inositol), suggesting their potential roles in root colonization [93–95].

Our analysis revealed that MAGs 1, 27, and 30, all assigned to phylum Pseudomonadota, possessed the necessary genes to produce the Flagellum-specific ATP synthase FliI enzyme, which is vital for the assembly and function of bacterial flagella [96, 97] (Figure 5, Table S7). This functionality is essential for motility, chemotaxis, adherence, and colonization. Additionally, we observed that MAGs 1 and 30 had a variety of genes (MAG 1: *chpA, flgH, flgI, flgG, fliP, fliG, fliM, fleQ, fleN, flil, pilT, pilM, pilG* and *pilH*; MAG 30: *flgB, flgH, flgC, flgG, fliP, flgE, filG, fliM, fliA, ppdD*, and *rfe*) potentially involved in the formation, regulation, and functioning of the bacterial flagellum [96, 98–100].

We observed that MAGs 9 (High GC gram+) and 27 (Pseudomonadota) could produce a long regulatory protein with the LuxR domain, commonly involved in bacterial Quorum Sensing and stress response systems (Figure 5, Table S7). Additionally, MAG 27 (Pseudomonadota) also had the potential to produce Acyl-homoserine lactones (AHLs), critical signaling molecules for N-butanoyl-L-homoserine lactone-binding transcriptional activation [101] (Figure 5, Table S7). In contrast, MAG 9 (High GC gram+) did not have the potential to produce AHLs signaling molecules because of the absence of a LuxI-encoding gene, suggesting that LuxR solos in MAG 9 (High GC gram+) might be involved in recognizing environmental stimuli and sensing host-derived signals [101, 102].

Additionally, we observed that MAGs 1 (Pseudomonadota) and 30 (Pseudomonadota) had a genomic potential to produce four essential products necessary for biofilm formation: PGA outer membrane secretin PgaA, PGA synthesis deacetylase PgaB, Biofilm PGA synthesis N-glycosyltransferase PgaC, and PGA synthesis auxiliary protein PgaD [49, 103] (Figure 5, Table S7). Since bacterial biofilms are multicellular communities formed by certain bacterial species to thrive in hostile environments, effective cell-to-cell communication through quorum sensing is essential for biofilm formation and maturation.

We then explored the potential for the MAGs to attach to the host roots [63, 104, 105]. We observed that MAG 1 (Pseudomonadota) could produce the outer membrane porin F (OprF) (Figure 5, Table S7) that can function as a root adhesion protein and play an essential role in root attachment [106]. Additionally, MAGs 9 (High GC gram+), 27 (Pseudomonadota), and 30 (Pseudomonadota) had various genes (MAG 30: hslO, amn, recA, ruvB, dacC; MAG 27: recA and ruvB; MAG 9: recA) that facilitate the attachment process [107]. We also observed that MAG 30 (Pseudomonadota) possessed genes pta and ompA (Figure 5, Table S7), which are associated with virulence and the invasion of host cells, thus contributing to intracellular colonization [108, 109].

Taken together, our gene- and genome-centric metagenome analyses corroborated with each other and with our previous hypothesis on the importance of *home-field advantage* in enhancing plant host resilience under drier environmental conditions. Microbial populations at both the community and genome levels displayed potential functions that could enable microbial survival and enhanced microbe-microbe-plant interactions under the drier environmental conditions in Hays. After analyzing the host and microbial functions separately, we next asked *which microbial functions could have the greatest impact on the plant host*.

### Ecotype-microbial interaction resulted in enhanced plant host resilience under drier conditions

In this study, our gene-centric analyses showed that plant-host stress-associated metabolites could enhance the recruitment of specific microorganisms, whereas genome-centric analyses identified the microbial population functions and plant-host interaction. With these insights, we next investigated the interaction between the *A. gerardii* ecotypes and their associated rhizobiome functions to determine which resultant interactions could enhance plant host drought resilience. Our DIABLO network analysis (cutoff=0.83) showed that MAG 34 (Thermoproteota) correlated negatively with host-derived peptide cyclo-(Gly-Phe) and trimethyllysine (TML) and positively with 3-amino-2-naphthoic acid, DL-Tryptophan, and 3,5-dihydro-2H-furo[3,2-C]quinolin-4-one (Figure 6A). Further, five MAGs (MAG 3 - Thermoproteota, MAG 4 - Actinobacteriota, MAG 5 - Desulfobacterota, MAG 10 - Desulfobacterota, and MAG 15 - Thermoproteota) positively correlated with Trimethyllysine (TML; Figure 6A). Interestingly, these five MAGs were detected more frequently in Hays than in Carbondale (Figure 6B, Figure 6B), and were also detected at higher levels in association with Dry ecotype in Hays. TML is important in plant metabolism, serving as a precursor to carnitine, a compound crucial for various metabolic and physiological functions [110, 111]. Similar to mammals and fungi, plant carnitine production begins with the breakdown of lysine-containing proteins, which then go through lysosomal or proteasomal protein degradation in a four-step pathway [110, 112]. Carnitine is essential for transporting fatty acids into the mitochondria for β-oxidation, which produces energy essential for various metabolic processes, particularly under abiotic stress [110]. In addition, one of the most common forms of carnitine - L-carnitine, serves multiple functions in plants, including detoxification, antioxidant activity, regulation of oxidative stress, lipid metabolism, stress signaling, and responses to oxidative damage during drought and high salinity [110, 111, 113]. Under drought stress, TML production and its conversion to carnitine could support the plant’s ability to manage oxidative stress and maintain cellular functions during water limitation. Despite the critical and diverse roles of TML in plants, our understanding of the role of TML is limited. We hypothesize that the lower TML levels produced by the Dry ecotype in Hays demonstrate the lower stress levels experienced by the Dry ecotype in its native (drier) environment. These findings support our two key hypotheses: 1) Dry ecotype in its native Hays environment experiences lower stress levels, leading to reduced production of stress-related metabolites, aligning with the “*home-field advantage*” hypothesis; 2) the positive association between the five MAGs and host produced TML suggest that these MAGs may contribute to alleviating plant stress or potentially provide a priming effect for the plant host. Although the lower TML levels were associated with the Dry ecotype in our network analyses, these differences were not statistically significant (Kruskal-Wallis: chi-squared = 1.64, df = 1, p = 0.200) (Figure 6C). We observed that this variability was driven by a population effect where the Dry populations (CDB and WEB) had significantly lower levels of TML compared to the Wet (DES and FUL) populations (Kruskal-Wallis: chi-squared = 8.44, df = 3, p = 0.038) (Figure 6D). Although the effect of population genetic diversity on metabolic profiling has been documented by others [114], this was the first instance in our analyses where it became evident [6]. These observations highlight the complexity of metabolic responses within different populations and suggest that factors such as local adaptation or genetic diversity might significantly influence the production of metabolites like TML. This underlines the importance of considering even finer population-specific resolution when studying host metabolic traits. With insights from the plant ecotypes, we asked next *what could be the interaction between the plant host and these 5 MAGs that would result in higher production of TML by the Dry ecotype in a drier environment (Hays)*.

**Figure 6.**
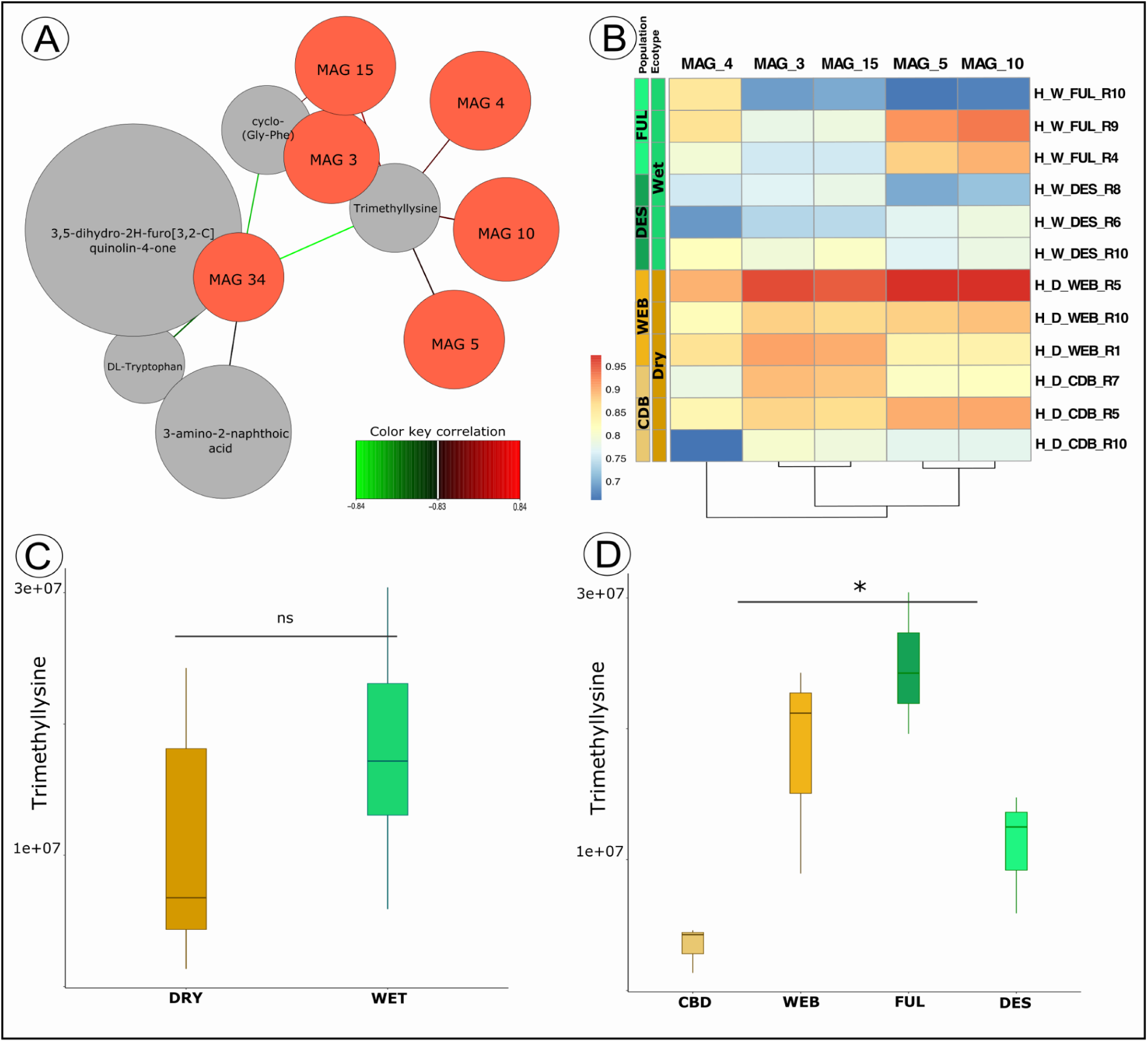
Plant host-derived metabolite, Trimethyllysine (TML) positively correlated with five MAGs and was highly associated with Dry ecotype populations in Hays. (A) Network analysis of plant-host metabolite (Grey) couples with MAGs detection levels (Red). A correlation cutoff of 0.83 was used to visualize only strong associations, with (Green) lines indicating negative correlations and (Red) lines marking a positive correlation of MAGs with the host metabolites. (B) The detection heatmap with rows corresponding to rhizosphere samples collected in Hays and columns corresponding to the five MAGs of interest. (C) The detection of the TML among Dry and Wet ecotypes in Hays. Not significant p-value marked by “ns.” (D) The detection of TML among Dry and Wet populations in Hays. p-value marked asterisks * represent p < 0.05.

Through our DIABLO analysis, we aimed to gain deeper insights into how the five MAGs associated with dry site interacted with plant-derived TML, particularly how these interactions might enhance the dry ecotype resilience in drier conditions such as those in Hays. We suggest that the relationship between TML, plant performance, and microbial activity is complex and neither simple nor unidirectional. The positive correlation of lower TML and the five MAGs suggests that they play a vital role in alleviating stress, thus enabling the host to better cope with stressful conditions in Hays. Consistently, the ability of these MAGs to thrive in the drier, more stressful environment in Hays might facilitate a healthier rhizosphere through several indirect mechanisms, including 1) improved nutrient availability and mobilization, 2) production of plant growth-promoting compounds, and 3) modulation of stress responses.

One avenue that may result in positive plant outcomes through TML-microbe interactions could be attributed to the ability of our resolved MAGs to use TML as a nutrient source to synthesize compounds critical for microbial stress tolerance and energy metabolism [110, 111, 115]. The resolved MAGs could synthesize carnitine from TML to stabilize their cell structures, manage oxidative stress, and maintain energy production under drier conditions in Hays [116]. For example, we observed that MAGs 4 (High GC gram+), 5 (Desulfobacterota), and 10 (Desulfobacterota) could produce L-carnitine dehydratase/bile acid-inducible protein F, an enzyme dehydrating L-carnitine to form other metabolites, which, in turn, would potentially make them available to the plant in different nutrient forms (Figure 7). We observed that all five MAGs possessed a diverse array of enzymes that catalyze the conversion of various substrates into ammonia: MAG 3 (glutamate-ammonia ligase), MAG 4 (glutamate-ammonia ligase, carbamate kinase, NAD-specific glutamate dehydrogenase, glutamate dehydrogenase (NADP+), and nitrate reductase), MAG 5 (aspartate ammonia-lyase, glutamate dehydrogenase (NADP+), and nitrate reductase), MAG 10 (glutamate-ammonia ligase, ferredoxin-nitrite reductase, glutamate dehydrogenase (GDH), glutamate dehydrogenase (NADP+), and nitrate reductase), and MAG 15 (glutamate-ammonia ligase). The ability of these five MAGs to convert substrates to ammonia could mobilize bioavailable nitrogen for the host, and enhance amino acid (including lysine) synthesis. Besides the ability to metabolize TML and to subsequently indirectly contribute to plant host through improved nutrient availability, our resolved MAGs also possess the capacity to produce plant growth-promoting compounds. Three out of these five MAGs possessed various gene functions related to the production of plant growth secondary metabolites (MAG 4 - Arylsulfatase, MAG 5 - Cyclic beta-1,2-glucan synthase, Arylsulfatase, 2,5-diketo-D-gluconic acid reductase; MAG 10 - 2,5-diketo-D-gluconic acid reductase, Putative arylsulfatase regulatory protein) and signaling molecules (MAG 4 - Inositol-1-monophosphatase; Phosphatidate cytidylyltransferase; Probable serine/threonine-protein kinase pknK; Serine/threonine-protein kinase PknB, PknD, PknE and MAG 5 and 10 - Inositol-1-monophosphatase) that may promote plant root development, enhancing root depth and spread, as reported in [117, 118] (Figure 7). We also surmised that the five MAGs could interact with plant host-derived TML, and enhance the plant drought tolerance through the production of stress-related enzymes (MAG 3 - AqpZ, MAG 4 - AqpZ, ABC transporter, ATP-binding protein and permease protein, Glycerol uptake facilitator protein and transporter OpuD, Cold shock protein of CSP family; MAG 5 - AqpZ, Cold shock protein of CSP family, Ribosomal arrest protein RaiA / Cold shock protein of CSP family; MAG 10 - Cold shock protein of CSP family, Ribosomal arrest protein RaiA / Cold shock protein of CSP family) and antioxidants (MAG 3 - Manganese catalase; MAG 4 - Manganese catalase, Hydroxyacylglutathione hydrolase; 16S rRNA (uracil(1498)-N(3))-methyltransferase; MAG 5 - Uncharacterized monothiol glutaredoxin ycf64-like, N-methylhydantoinase A and B, stress proteins YciF and YciE, Glutathione synthetase, Glutathione reductase, Glutathione peroxidase and Thioredoxin peroxidase, Glutathione S-transferase family protein, Glutamate--cysteine ligase, Gamma-glutamyltranspeptidase and Glutathione hydrolase, Catalase KatE; MAG 10 - Hydroxyacylglutathione hydrolase, Glutathione reductase, Glutathione peroxidase and Thioredoxin peroxidase, Glutathione S-transferase; MAG15 - Superoxide dismutase [Mn], Manganese catalase) (Figure 7). The production of microbe-derived antioxidants from these MAGs could reduce oxidative damage in the plant host under the drier and more stressful conditions in Hays, thus improving plant growth and productivity.

**Figure 7.**
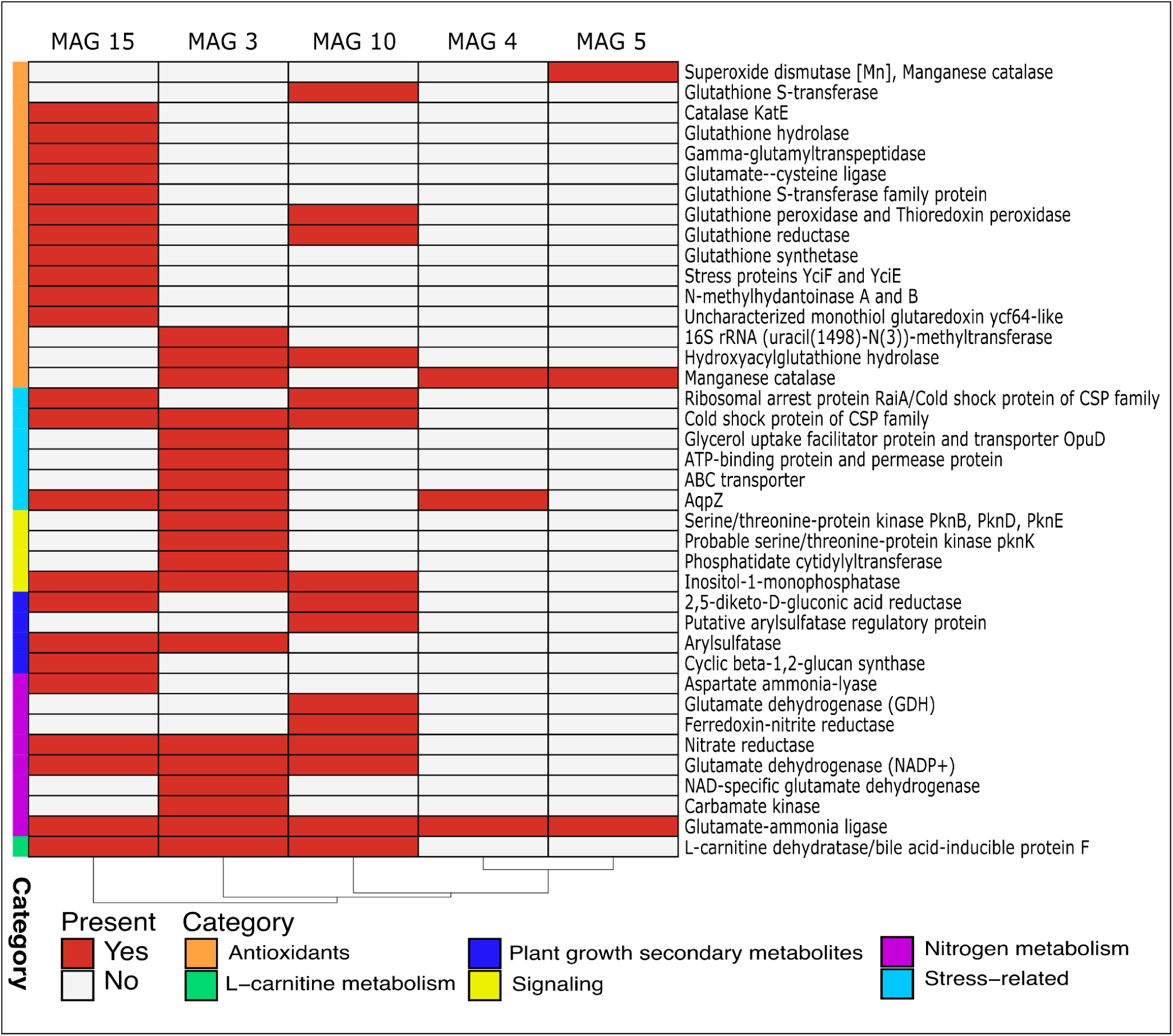
Genes/subsystems (subset) presence (red) and absence (grey) in the MAGs-of-interest. Rows correspond to the subset of genes/subsystems related to microbial mechanisms detected in five MAGs (columns) of interest, potentially promoting plant-host survival and environmental performance.

Additionally, TML is a precursor to trimethylamine N-oxide (TMAO) in the human system [119, 120]. Some research is starting to uncover TMAO’s potential in terrestrial plant systems, and its role in stress mitigation suggests a huge potential in supporting plants in marginal environments. While we did not find direct evidence of this conversion in the five MAGs of interest, we did detect trimethylamine-N-oxide reductase (TorZ) and trimethylamine-N-oxide reductase and associated c-type cytochrome (TorY) in MAG 27, suggesting its capacity to metabolize TML [121–123]. TMAO is increasingly recognized for its potential role in improving plant stress tolerance, especially in challenging environmental conditions, e.g., drought and heat [124, 125]. The microbial TMAO, and in our study, specifically MAG 27, could act as osmoprotectants. Microbial populations can metabolize TMAO to improve their resilience to osmotic stress [124–126], allowing them to maintain metabolic activity around plant roots under dry conditions such as those in Hays. Although known only in microbial populations, it is conceivable that TMAO could stabilize proteins and cellular structures in plant cells as well [124, 126, 127]. TMAO’s protein and enzyme stabilizing effects might also extend to antioxidant enzymes in the plant host [124, 126], protecting the plant ecotypes from oxidative stress associated with environmental stresses in Hays and helping them maintain cellular integrity and physiological functions under drier conditions.

In sum, the interactions between plant-derived TML, ecotypic microbial populations, and TML’s derivatives strongly suggest that there exists a deliberate plant control of the associated microbes through plant-produced metabolites. We understand that the interaction between the plant-derived TML and associated microbial constituents extends beyond these few MAGs present in our study. However, our results set the platform to motivate new lines of investigation as they provide a rational, mechanistic basis to offer deeper insights into complex plant-microbe interactions. These findings further support the “*home-field advantage*” hypothesis, showing that the plant actively produces metabolites to recruit local microbial populations. These functional capabilities ultimately contribute to the host’s ability to thrive in stressful environments. The variability in the TML production among plant ecotypes and populations further highlighted the complexity of the plant metabolic responses shaped by genetic diversity and the plant’s local adaptations. Further research on TML and related processes is needed. Our current research deepens our understanding of the factors that drive rhizobiome recruitment and offers insights into how rhizobiome communities contribute to ecotype responses to climate change.

## Conclusions

This study highlights the intricate interplay between plant-derived metabolites and rhizobiome community functions to enhance *A. gerardii* drought tolerance. Our data indicate that the interaction between plant host-derived trimethyllysine (TML) and the microbial capacity for chemotaxis, biofilm formation, and root attachment are more characteristic mechanisms of plant stress resilience than previously appreciated.

Our findings underscore how the functional potential of the rhizobiome communities is influenced by plant metabolites. Our work expands on the “*home-field advantage*” hypothesis, demonstrating that plants adapted to drier environments recruit microbial populations enriched in pathways that mitigate abiotic stress. Specifically, our data show that plant-derived metabolite TML correlates with microbial populations that enhance plant resilience through nutrient cycling, antioxidant production, and stress-signaling modulation.

This study and our methodological framework, combining host metabolomics with microbial gene-centric and genome-centric analyses, provide a blueprint for future studies exploring plant-rhizobiome interactions. By linking host metabolomics to microbial genomic potential, we are able to systematically identify microbial pathways that are critical for stress tolerance. Our approach can also be applied to other plant systems to investigate stress-response mechanisms or to engineer microbiomes to enhance agricultural productivity. This study advocates for a shift from taxonomic focus to functional rhizobiome characterizations. Future work should explore how microbial metabolites may contribute to host resilience and expand our understanding of metabolite-mediated microbial recruitment.

While our study identifies important host metabolites and microbial functions, some questions remain unanswered, specifically the signaling pathways through which TML mediates host-rhizobiome interactions. Future work should integrate metatranscriptomics to capture the real-time dynamics of these interactions. In summary, our work demonstrates that plant metabolite-microbial function interactions are a cornerstone of rhizosphere assembly and plant resilience to environmental drought stress. By leveraging these insights, we can design microbiome-based solutions to improve plant performance under climate change, paving the way for successful conservation efforts and sustainable agricultural practices.

## Methods

Detailed sampling methods are described in Kazarina et al. [6] and briefly summarized here. Following our previous work [6], we asked what plant-associated and microbial-associated functions in the plant-host rhizobiome interaction contribute to the “home-field” advantage and could potentially enhance *Andropogon gerardii* drought stress resilience.

### Selection of rhizosphere samples and shotgun metagenomics

We selected 24 *A. gerardii* rhizosphere samples to gain deeper insights into microbial functions that could potentially enhance plant resilience and responses to microbial interactions during drought stress (Table S1). Samples were chosen partially based on the results of Kazarina et al. [6] to better understand the host-microbe interactions and consisted of samples from two reciprocal gardens located in two precipitation extremes, (H - Hays, Kansas (580 mm/year) and C - Carbonate, Illinois (1,167 mm/year), planted with two regional ecotypes (Dry and Wet) and 4 populations (Dry: CDB, WEB; Wet: DES, FUL) (2 locations x 2 ecotypes x 2 population x 3 samples) (Table S1). We extracted total gDNA from 0.150 g of roots and rhizosphere soil from all 24 samples using the E.Z.N.A Soil DNA Kit (Omega Bio-Tek, Inc., Norcross, GA, USA), following the manufacturer protocol with slight modifications. Roots were shaken to remove non-adhering soil and weighed for the DNA extractions. Any soil that remained attached to the roots was considered rhizosphere soil [128, 129]. We mechanically lysed the samples on a Qiagen TissueLyser II (Qiagen, Hilden, Germany) using glass beads, for 10 mins at 20 rev/s prior to any downstream DNA extraction steps. The mechanical lysis never fully homogenized the roots; therefore, our samples represented largely rhizosphere microorganisms, potentially excluding many endorhizosphere organisms. The extracted gDNA was eluted to 100 µL final volume. The DNA yield and concentration were measured using a Nanodrop and a Qubit™ dsDNA BR Assay Kit. Extracted DNA was kept at −20°C until shotgun metagenome sequencing. Extracted gDNA was used to construct DNA libraries using Illumina’s Nextera XT Index Kit v2 kit. The products were visualized on an Agilent Tapestation 4200 and size-selected using the BluePippin. The final library pool of 24 samples was quantified using the Kapa Biosystems qPCR protocol and sequenced on the Illumina NovaSeq S1 flow cell chip in a 2 × 150bp paired-end sequencing run.

### Bioinformatic workflow of gene-centric metagenomic analyses

We used biobakery meta-omics analysis environment [130] to taxonomically and functionally profile the microbial communities in the metagenome samples. The taxonomic profiling was assessed using the MetaPhlAn v.4.1.1 [131] with CHOCOPhlAn v3.0.14 [132] database. For functional profiling, we used HUMAnN3 v.3.9 [132] pipeline with MetaPhlAn v.4.1.1, Bowtie2 version 2.2.6 [133], DIAMOND v 2.0.15 [134], and MinPath v.1.5 [135]. HUMAnN3 pipeline was performed with the default parameters.

### Bioinformatic workflow of MAGs assembly and genome-centric metagenomic analysis

We used automated metagenomics bioinformatics workflows implemented by the programs ‘anvi-run-workflow’ [136] in anvi’o [137, 138]. Anvio’s workflows implement bioinformatic tasks, including short-read quality filtering, assembly, gene calling, functional annotation, hidden Markov model search [139], metagenomic read-recruitment, metagenomic binning [140], pangenomics, and phylogenomics. Workflows use Snakemake [141], and a tutorial can be found at the URL https://merenlab.org/2018/07/09/anvio-snakemake-workflows/.

We used ‘iu-filer-quality-minoche’ to process the short metagenomic reads, implemented in illumina-utils v2.11 [142], and removed low-quality reads according to the criteria outlined in Minoche et al. [143]. We used MEGAHIT v1.2.9 [144] to co-assemble quality-filtered short reads into longer contiguous sequences (contigs). Due to the high sample heterogeneity, we split the metagenomes into 4 even experimental groups based on location and plant ecotypes (location-ecotypes: H-Dry, H-Wet, C-Dry, and C-Wet) for the co-assembly. We used the following strategies to process contigs: (1) ‛anvi-gen-contigs-databasè on contigs to compute k-mer frequencies, and identify open reading frames (ORFs) using Prodigal v2.6.3 [145]; (2) ‘anvi-run-hmms’ to identify sets of bacterial [146] and archaeal [147] single-copy core genes using HMMER v.3.2.1 [148]; (3) ‛anvi-run-ncbi-cogs‛ to annotate ORFs with functions from the NCBI’s Clusters of Orthologous Groups (COGs) [149], and (4) ‘anvi-run-kegg-kofams’ to annotate ORFs from KOfam HMM databases of KEGG orthologs [150, 151].

We recruited metagenomic short reads to contigs using Bowtie2 v2.3.5 [133], and converted the resulting SAM files to BAM files using samtools v1.9 [152]. We profiled the BAM files using ‘anvi-profile’ with a minimum contig length of 1000 bp to eliminate shorter sequences and minimize noise. Each metagenome was then profiled to store contig coverages into single anvi’o profile databases, and we used ‛anvi-merge’ to combine all profiles into an anvi’o merged profile for downstream visualization, binning, and statistical analyses. We used ‘anvi-cluster-contigs’ to group contigs into initial bins using CONCOCT v1.1.0 [153], and used ‘anvi-refine’ to manually curate the bins based on tetranucleotide frequency and different coverage across the samples. We marked bins that were more than 70% complete and less than 10% redundant as metagenome-assembled genomes (MAGs). We used the “detection metric” to assess the presence of MAGs in a given sample, and considered a MAG as detected in a metagenome if the detection value was > 0.25, which is an appropriate cutoff to eliminate false-positive signals in read recruitment results, for its genome.

### Phylogenomic analysis and functional potential of MAGs

We uploaded and annotated MAGs on the Bacteria and Viral Bioinformatics Resource Center (BV-BRC) platform [154]. We used the similar genomes finder function in BV-BRC to download 11 similar genomes for each MAG (a total of 34 MAGs and 324 similar genomes). We used a collection of six bacterial ribosomal gene alignment to construct the phylogenetic tree: Ribosomal_L1, Ribosomal_L2, Ribosomal_L3, Ribosomal_L4, Ribosomal_L5, Ribosomal_L6. We explored these sequences across all the assembled draft genomes and similar genomes using anvi’o program ‘anvi-get-sequences-for-hmm-hits’, and aligned the sequences of each gene using MUSCLE v3.8. [155]. We then used IQ-TREE v1.6.12 [156] with the ‘WAG’ general matrix model [157] and 1,000 ultrafast bootstrap to compute maximum likelihood phylogenetic trees for each alignment. Finally, we used anvi’o v7.1 program ‘anvi-interactive’ program to visualize the constructed phylogenetic trees and associated metadata. Following the confirmation of identity, we utilized Subsystems genes [158] and pathway functions in the KEGG database [151], to compare the potential functions between MAGs.

### Metabolomic profiling of plant host rhizosphere

#### Sample preparation

Samples selected for plant host metabolite profiling were collected during early Fall 2023 within one week of each sampling time (C - September 03, 2023, H - September 13, 2023). Roots were picked from the selected samples, and shaken to remove any soil; the soil that remained on the roots was considered rhizospheric soil. Rhizosphere soil was separated from the roots, and a 20 g soil sample was aliquoted into a 50 mL centrifuge tube. We added 25 mL of deionized water to each of the 50 mL tubes and mixed the contents by shaking for 3 hours at room temperature. Samples were then centrifuged for 5 mins at 8000 rpm. The supernatant was transferred to the new 50 mL tube, and kept at −80°C until metabolomic analyses (MetWare® Bio, MA, USA).

#### Sample extraction and chromatography-mass spectrometry

An aliquot of each sample (9 mL) was placed at 4°C overnight, and then vacuum freeze-dried. After freeze-drying, 70% methanolic internal standard extract was added at a ratio of 30 times the concentration. Samples were vortexed for 15 mins, and sonicated in an ice water bath for 10 mins. The extracts were centrifuged at 12,000 rpm at 4°C for 3 mins, and filtered using a 0.22 μm filter before liquid chromatography and mass spectrometry. Ultra-performance liquid chromatography (UPLC) (ExionLC™ AD) and tandem mass spectrometry (MS/MS) (Applied Biosystems QTRAP 6500) were used for data acquisition. Liquid phase condition consisted of (1): liquid phase chromatographic column: Agilent SB-C18 1.8 µm, 2.1 mm × 100 mm; (2) Mobile phase: A phase—ultrapure water (0.1% formic acid added), B phase—acetonitrile (0.1% formic acid added); (3) elution gradient: the proportion of B phase was 5%, within 9.00 min, the proportion of B phase increased linearly to 95% and remained at 95% for 1.00 min, 10.00–11.10 mins, the proportion of B phase decreased to 5% and balanced at 5% up to 14.00 mins; (4) flow rate: 0.35 mL/min; (5) column temperature: 40°C; and (6) injection volume: 2 μL.

Linear Ion TRAP Mass Spectrometer (Q TRAP) (AB6500 Q TRAP UPLC/MS/MS system) was used to obtain both mass spectrum scans using both Linear Ion TRAP Mass Spectrometer modes. Operating parameters of ESI source were as follows: ion source, turbine spray; source temperature 550°C; Ion spray voltage (IS) 5500 V (positive ion mode) / −4500 V (negative ion mode); The ion source gas I (GSI), gas II (GSII) and curtain gas (CUR) were set to 50, 60 and 25.0 psi respectively, and the collision-induced ionization parameter was set to high. The multiple reaction monitoring (MRM) mode was used for the QQQ scan and collision gas (nitrogen) was set to medium. DP and CE of each MRM ion pair were completed by further DP and CE optimization, and a specific set of MRM ion pairs was monitored based on the eluted metabolites at each period.

Secondary metabolites were identified based on their secondary spectrum information. Signals from isotopes, and repeated signals containing cations such as K+, Na+, and NH ^+^ were excluded during the qualitative analyses. Metabolites were quantified by triple quadrupole mass spectrometry with multiple reaction monitoring (MRM). In MRM mode, the first quadrupole screened precursor ions specifically for the target compound, excluding ions of other molecular weights. After ionization induced by the impact chamber, the precursor ion was fragmented, and a characteristic fragment ion was selected through the third quadrupole, and excluded the interference of other non-target ions.

#### Data preprocessing

Missing values in the acquired raw data were filled using 1/5 of the minimum value of each metabolite, and then the Coefficient of Variation (CV) value of the quality control (QC) sample was calculated, and the metabolites with a CV value less than 0.3 were removed from the further analysis. The CV signifies the relationship between the standard deviation and the mean in the original dataset and was used as an indicator of data dispersion. The Empirical Cumulative Distribution Function (ECDF) was used to analyze the frequency of compound CVs falling below a specified reference value. A composite QC specimen was prepared from the mixture of extracts from all samples to assess the reproducibility of the entire metabolomics workflow. Throughout data acquisition, a single quality control sample was systematically incorporated for every 10 test samples.

The mass spectrum data was processed using Analyst 1.6.3 (AB Sciex, Framingham, Massachusetts, USA). The MRM metabolite multi-peak detection diagram depicting the compounds identified within the sample was generated. Each peak in the mass spectrum is color-coded to signify the detection of a specific metabolite. The characteristic ions of each compound were selected by triple quadrupole, and their signal intensities (CPS - Counts Per Second) were measured. Subsequently, the mass spectrometry data were analyzed through MultiQuant software, facilitating the integration and correction of chromatographic peaks. The area of each chromatographic peak represents the relative abundance of the corresponding compound. The correction of mass spectrum peaks for each metabolite across various samples was conducted by considering retention time and peak distribution data to ensure the precision of both qualitative and quantitative analyses.

We used the vegan package [159] in R Studio to estimate the pairwise Bray–Curtis distances to compare the relative metabolic content of metabolites among the different samples. We then used the ggplot2 package [160] to visualize these data using non-metric multidimensional scaling (NMDS) ordinations.

### Statistical analysis

We (1) first identified key omics variables for the integration process; and then (2) maximized the common and correlated information between the datasets, and finally (3) visualized the results to identify relevant microbial metagenomics and host metabolomics correlations. We used Analysis of Variance (ANOVA) to test for main and interactive effects among groups. Following the overall ANOVA, we used pairwise comparisons using Pairwise Wilcoxon Tests with the false discovery rate (FDR) method to correct for multiple comparisons to identify factors that were driving the significant effects. P-values of less than 0.05 were considered statistically significant. Statistical analyses were performed in RStudio [161]. We used the program - Data Integration Analysis for Biomarker discovery using Latent cOmponents (DIABLO) to identify the key correlations between host metabolomics and metagenomic datasets (MAGs, MetaPhlAn, and HUMAnN) [162–165]

## Declarations

### Ethics approval and consent to participate

Not applicable.

### Consent for publication

Not applicable.

### Availability of data and materials

The datasets generated and/or analyzed during the current study are available in the Sequence Read Archive repository under NCBI BioProject PRJNA1181896. All other analyzed data files and codes are accessible at GitHub (https://github.com/SonnyTMLee/Interaction-of-plant-derived-metabolites-and-rhizobiome-functi ons-enhances-drought-stress-tolerance).

### Authors’ contributions

A.J., L.J., and S.T.M.L. designed the study. L.J. designed the reciprocal gardens experimental design, and A.J., L.J., E.H., and S.T.M.L. maintained the sites. A.K., S.S., S.P., and S.T.M.L performed sample collection. S.P., A.K., B.A., L.R., and S.S. extracted DNA and performed quality analysis with Nanodrop and Qubit for the metagenomics analysis. A.K., B.A., and L.R. prepared samples and processed data for the metabolomics analysis. A.K. and S.T.M.L. performed gene-, genomic-centric, and metabolomics data analyses. A.K. and S.T.M.L wrote the manuscript, and prepared figures, and files. A.J. and L.J. aided in manuscript and figure revisions. S.T.M.L., A.J., and L.J. acquired funding for this study. All authors read, contributed to the manuscript revision, and approved the submitted version.

## Acknowledgments

We would like to thank all of those who assisted in sampling and data acquisitions, especially Ricky Blakesleay, Jack Sytsma, Shiva Thapa, and Trey Summers. We would like to acknowledge Clark Bloomer and his team at the Genome Sequencing Facility at the University of Kansas Medical Center (https://www.kumc.edu/research/genome-sequencing-facility.html) for the technical support with the shotgun metagenomic sequencing. We appreciate Jeff Chu at MetWare Bio (https://www.metwarebio.com/) for the services and help with the metabolomics data generation. In addition, we are grateful for the help of Tanner Richie with the discussion on bioinformatics and data analysis.

## Authors information

X: @sonnytmlee (Sonny T.M. Lee); @annaKazarina2 (Anna Kazarina); Bluesky: @sonnytmlee..bsky.social (Sonny T.M. Lee); @akazarina.bsky.social (Anna Kazarina)

## Corresponding author

Sonny T.M. Lee -

## Tables

Table S1. The description of the *Andropogon gerardii* rhizosphere collection samples used for the metagenome and metabolomics analysis

Table S2. Metabolomic analysis results include the overall analysis, analysis by location (Carbondale and Hays), ecotype (Dry and Wet), population (DES, FUL, WEB, and CDB), and location and ecotype interactions.

Table S3. Gene-centric community profiling (MetaPhlAn), including the overall output and the subset resolved to genus level analysis by location (Carbondale and Hays), ecotype (Dry and Wet), and location and ecotype interactions.

Table S4. Gene-centric metabolic profiling (HUMAnN) including the overall output and analysis by location (Carbondale and Hays), ecotype (Dry and Wet), population (DES, FUL, WEB, and CDB), and location and ecotype interactions.

Table S5. Shotgun metagenome sequencing data obtained from the 24 rhizosphere samples used for the gene- and genome-centric analysis

Table S6. Description of the 34 MAGs with the ANVI’O taxonomic assignment.

Table S7. The subset of gene/subsystems with the function to promote plant-host growth or communicate with plant-host growth or other microbes observed in the 34 MAGs.

